# Description of *Protosticta sexcoloratus* sp. nov. (Odonata, Platystictidae) from the Western Ghats, India

**DOI:** 10.1101/2023.09.25.559450

**Authors:** Ayikkara Vivek Chandran, P K Muneer, M Madhavan, Subin K. Jose

## Abstract

*Protosticta* Selys, 1885 is a speciose genus of damselflies distributed in the tropical and subtropical forests of Asia. During an ongoing study to document the odonate diversity of the Western Ghats, we came across a colony of *Protosticta* species in Wayanad, Kerala, that appeared different from all other species hitherto described. We describe this population as a new species after detailed morphological comparison with closely similar species occurring in the region.

## Introduction

Platystictidae Kennedy, 1920 is a family of damselflies that are commonly known as shadowdamsels, distributed throughout Asia, Central America and South America (Paulson 2009). Most members of this family are forest dwellers and breed in small streams. *Protosticta* Selys, 1885 is a genus of shade-loving, slender representatives of this family found in the tropical and subtropical forests of south and southeast Asia. These damselflies inhabit the undergrowth of dense jungle, usually near hill streams. They are dark, with long, thin abdomen and structurally complex caudal appendages. They have relatively small interspecific differences, mostly in their prothorax and caudal appendages. It is a diverse group with 55 species described till date (Paulson et al. 2023). The Western Ghats of India has 15 described *Protosticta* species, many of which were discovered only recently (Joshi et al. 2020, Sadasivan et al. 2022, Vijayakumaran et al. 2022, Payra et al. 2023). Most of these damselflies have highly restricted distribution range within the Western Ghats, and are hence, of conservation concern.

In our endeavour to investigate the odonate diversity of the Western Ghats, we came across a population of *Protosticta* in Vellarimala, Wayanad, which had the following remarkable characters observable in the field: relatively small size for members of the group, considerable difference in coloration between the sexes which were of the same size, pale purple prothorax in male with black markings (Figures 1 and 2). We collected some individuals of this population and performed a detailed morphological comparison with published descriptions of all *Protosticta* species of the Western Ghats and other closely similar species collected from Wayanad, which allowed us to describe the population from Vellarimala as a species new to science.

**Figure 1:**
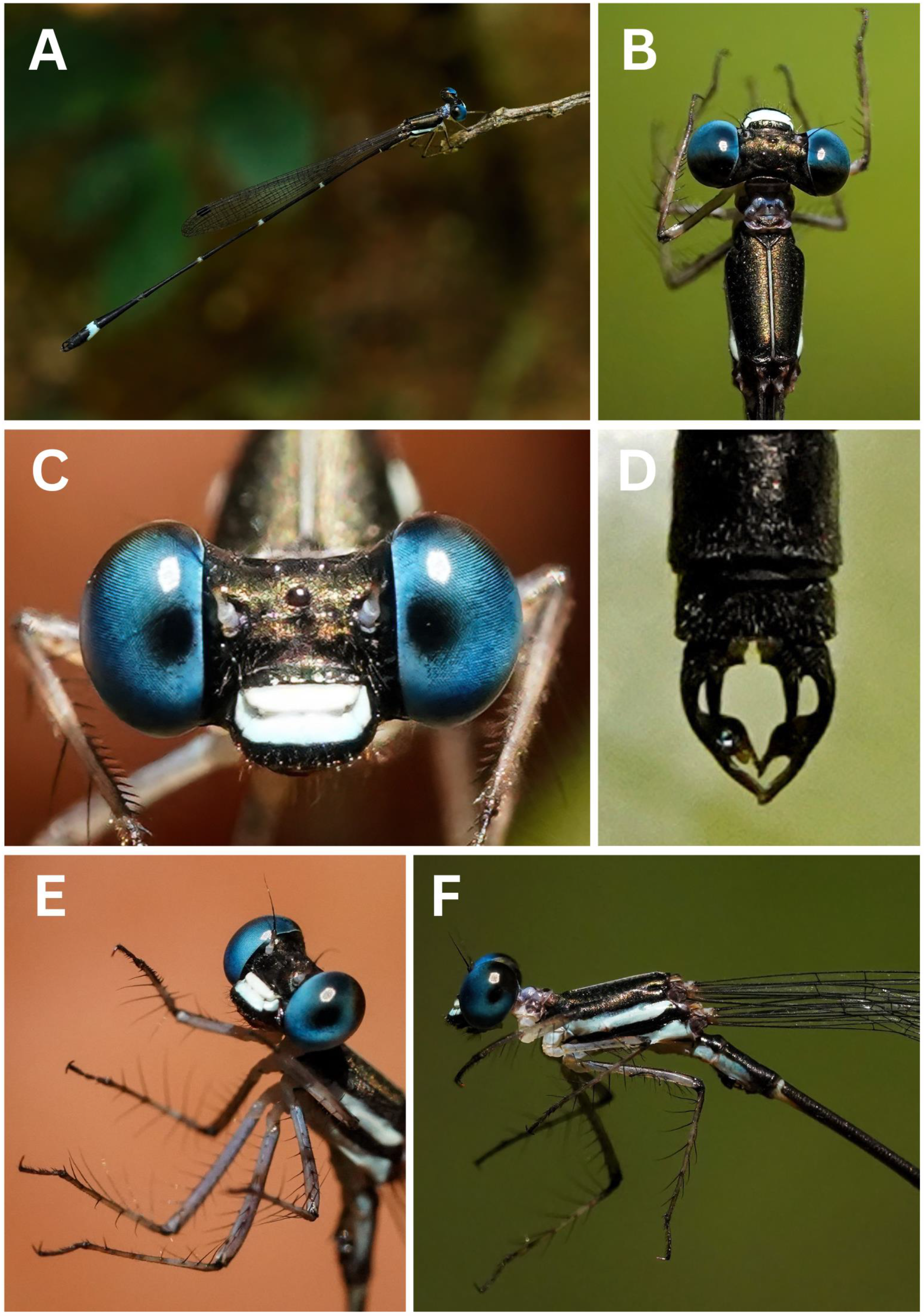
Live photographs of *Protosticta sexcoloratus* sp. nov. male, A- in the habitat, B- dorsal view of head and thorax, C- face, D- caudal appendages, E- legs, F- lateral view of head and thorax.

**Figure 2:**
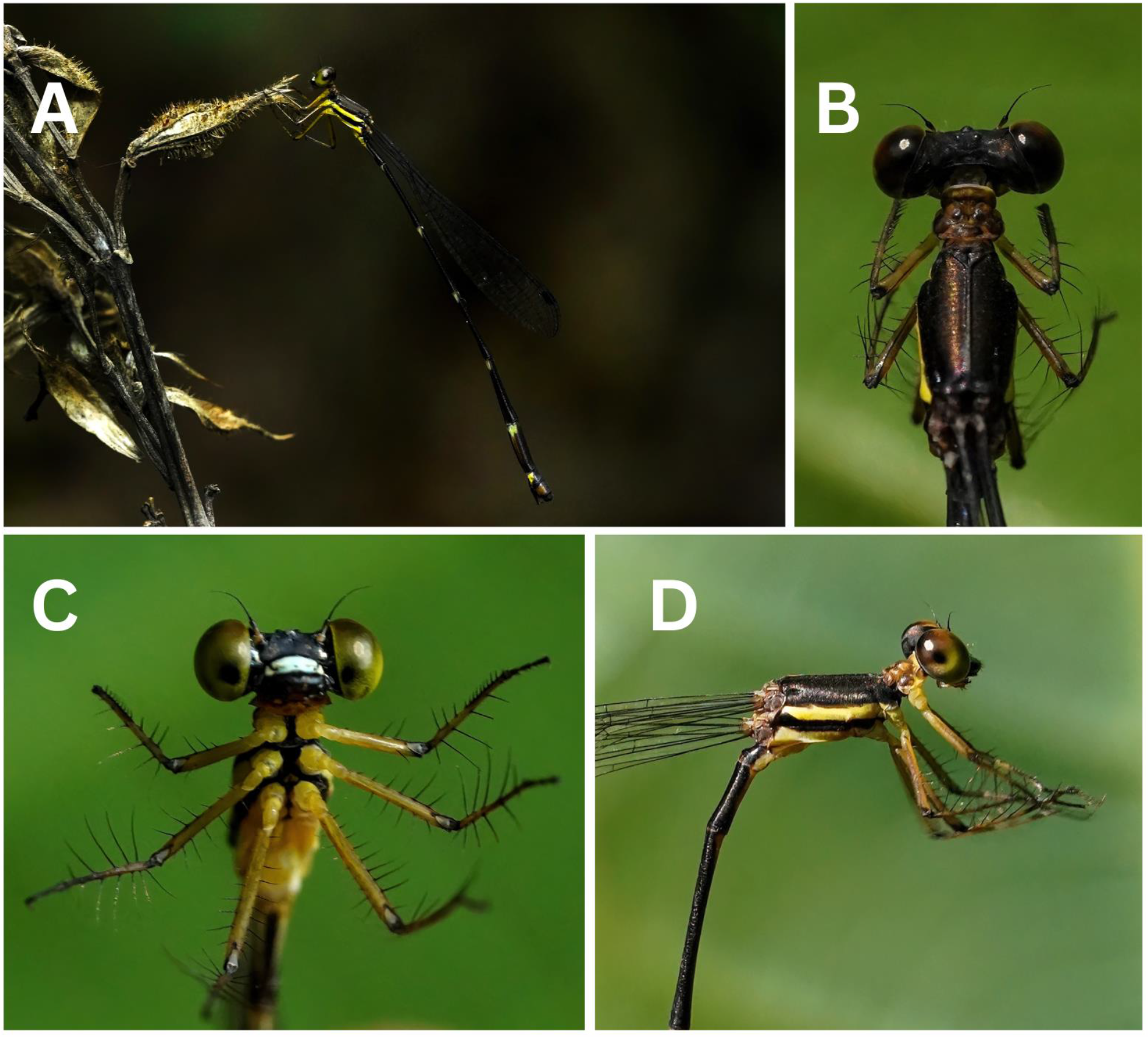
Live photographs of *Protosticta sexcoloratus* sp. nov. female, A- in the habitat, B- dorsal view of head and thorax, C- head, legs and venter of thorax, D- lateral view of head and thorax.

## Material and methods

*Protosticta* species were collected using a butterfly net, from three locations in Wayanad district of Kerala state, India (Figure 3). Three male and two female specimens of *Protosticta sexcoloratus* sp. nov. were collected from Vellarimala, Meppadi Forest Range, Wayanad (11.468920° N, 76.148280° E, 1352 m above MSL). One of the male specimens was put in absolute ethanol and the remaining specimens were preserved dry. A male specimen each of *Protosticta mortoni* Fraser, 1924 and *Protosticta gravelyi* Laidlaw, 1915 were collected from Brahmagiri (11.915740° N, 75.997700° E, 923 m above MSL) and Pulayankolli (11.903748° N, 76.030068° E, 835 m above MSL) respectively, for comparative study. These were preserved in absolute ethanol after taking detailed photographs. Field photographs of *P. sexcoloratus* sp. nov. and *P. mortoni* were taken using a Sony a7III mirrorless digital camera, Tamron 150-500 mm telephoto lens and Sony 90 mm macro lens. Field photographs of *P. gravelyi* were taken using Nikon Z6II mirrorless digital camera and Nikkor Z MC 105 mm macro lens. For taking photographs of preserved specimens, a Raynox DCR 250 supermacro lens was also used, coupled with the Sony equipment. The specimens in absolute ethanol were dissected under a stereomicroscope (SkiHi TDLED-1005, India) for studying their genital ligulae. All measurements were made using a digital Vernier calliper (ZHART CT-ZT-VERNIER). Diagrams of genital ligulae and caudal appendages were made using the Android application Sketchbook. Descriptive terminology follows Garrison et al. (2010). Quantum GIS (QGIS) version 3.22 and Google Earth Pro were used to create a map of the study area.

**Figure 3:**
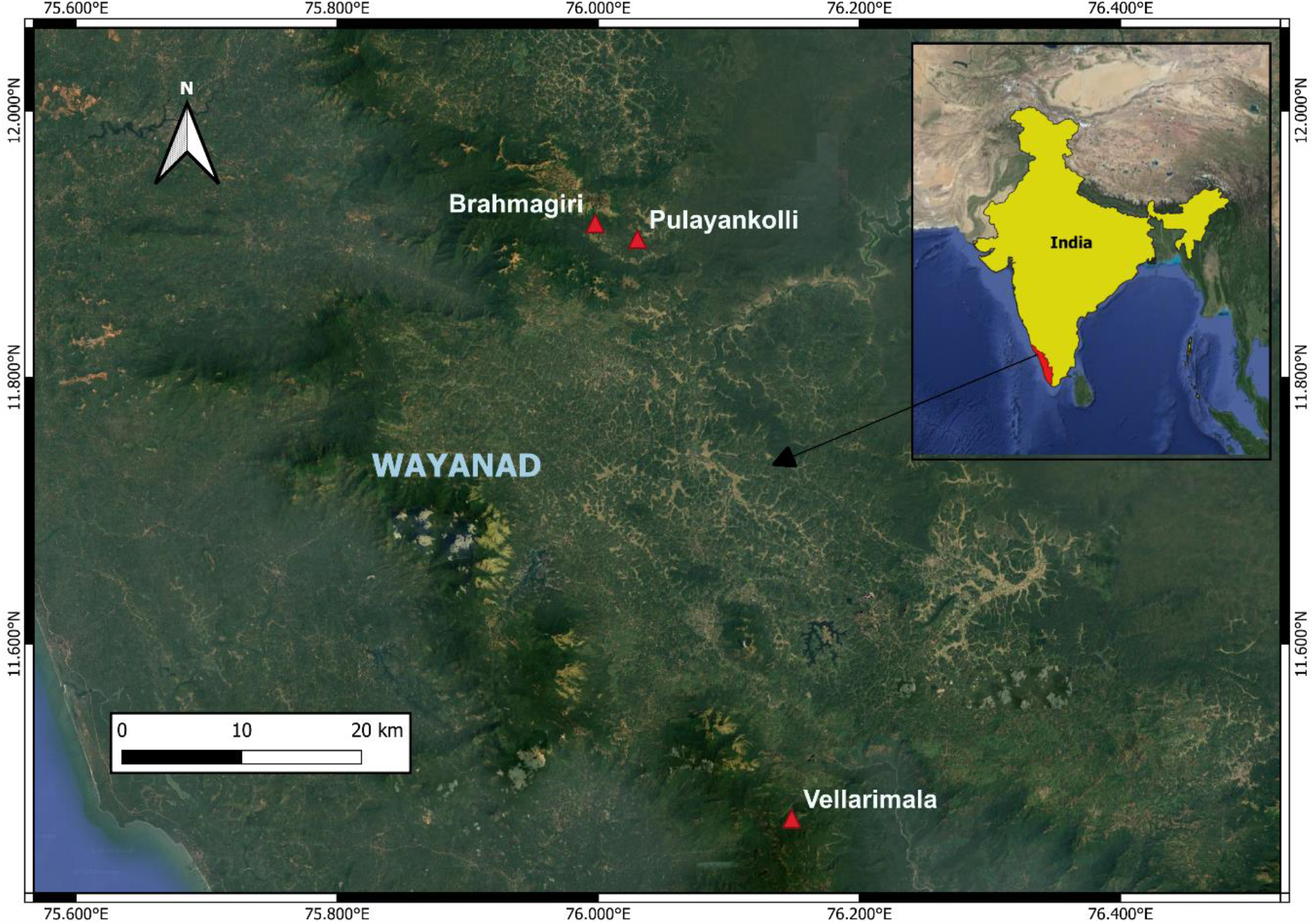
Locations of collection of *Protosticta sexcoloratus* sp. nov. (Vellarimala), *P. mortoni* (Brahmagiri) and *P. gravelyi* (Pulayankolli).

Type specimens and preserved genital ligulae are now in the insect collections of the Department of Geology and Environmental Science, Christ College (Autonomous), Irinjalakuda (ICDGE). The holotype will be deposited in the National Designated Repository for Fauna, Western Ghat Regional Centre, Zoological Survey of India (WGRC, ZSI) in due course. The following abbreviations are used in this study: FW= Fore wing, HW= Hind wing, Ab= Anal bridge, Ax= Antenodal nervures, Px= Postnodal nervures, Pt= Pterostigma, S1-S10= first to tenth abdominal segments, TL= Total length, AB= Abdomen, CA= Caudal appendages. Specimen label data are quoted verbatim.

### Systematic accounts

Order Odonata Fabricius, 1793

Suborder Zygoptera Selys, 1854

Superfamily Platystictoidea Kennedy, 1920

Family Platystictidae Kennedy, 1920

**Genus Protosticta** Selys, 1885

Type species: *Protosticta gravelyi* Laidlaw, 1915

***Protosticta sexcoloratus*** Chandran, Muneer, Madhavan and Jose sp. nov.

urn:lsid:zoobank.org:pub:71C8984A-D4C6-4C46-A664-D5FE77E85D3E

*Type material*. Holotype. ♂, labelled as “Holotype *Protosticta sexcoloratus* Chandran, Muneer, Madhavan, Jose, India, Kerala, Wayanad, Vellarimala [11.468920° N, 76.148280° E], elevation 1352 m above MSL, 21 v 2023, Coll. A. Vivek Chandran” (WGRC, ZSI).

Paratypes. 2 ♂ and 2 ♀, data as holotype (ICDGE).

*Diagnosis*. The most striking feature of *Protosticta sexcoloratus* sp. nov. is that unlike in most *Protosticta* species, the male and female are colored very differently. The following differential diagnosis of male *Protosticta sexcoloratus* sp. nov. is based on recently published descriptions and keys (Joshi et al. 2020, Sadasivan et al. 2022, Vijayakumaran et al. 2022, Payra et al. 2023).

*Protosticta sexcoloratus* sp. nov. can be distinguished from all other *Protosticta* species except *Protosticta hearseyi* Fraser, 1922 (Figure 4) by the fine blue middorsal carina of the male. Also, the sexes are of equal length in these two species. It can be told apart from *P. hearseyi* by the pale purple prothorax marked with black (fully pale blue prothorax in *P. hearseyi*), unmarked S9 (S9 with a ‘crescent and star’ marking in *P. hearseyi*) and rounded tips of cerci (cerci squared at tips in *P. hearseyi*).

**Figure 4:**
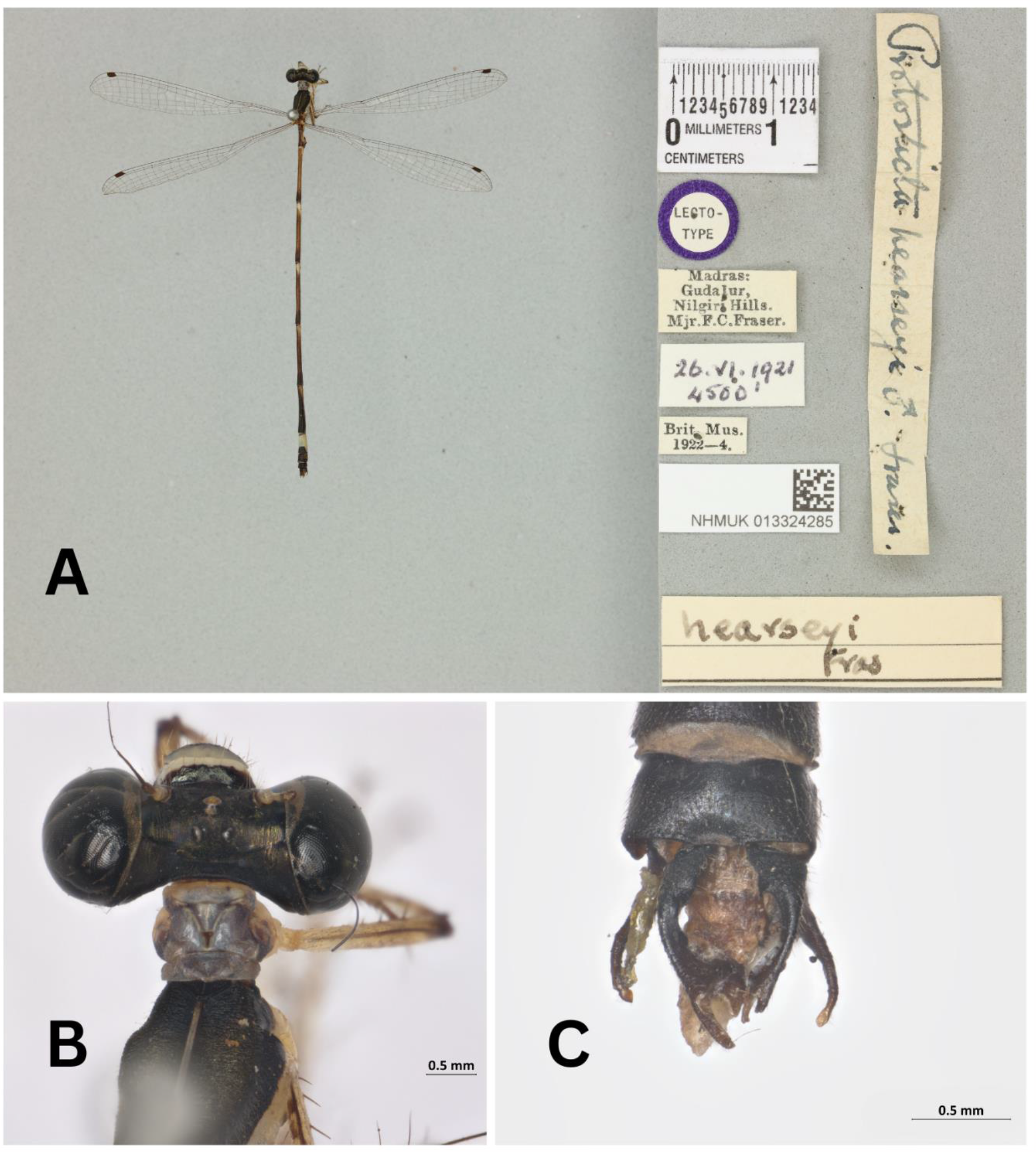
Images of *Protosticta hearseyi* lectotype from British Natural History Museum, A- habitus, B- prothorax, C- dorsal view of caudal appendages.

The new species is much smaller (AB+CA < 35 mm) compared to the three large species found in the Western Ghats, namely, *Protosticta antelopoides* Fraser, 1931, *P. francyi* Sadasivan, Vibhu, Nair & Palot, 2022 and *P. ponmudiensis* Kiran, Kalesh & Kunte, 2022 (AB+CA > 50 mm). It also lacks spines on the posterior lobe of prothorax. Also, in the new species, paraprocts are not bifid and have prominent sub-basal spines.

The new species has caudal appendages that are closely similar to those of *Protosticta gravelyi* Laidlaw, 1915 (Figure 5) and *P. mortoni* Fraser, 1924 (Figure 6), but differs in the shape of the sub-basal, inwardly directed spine on cerci (Figure 7). It also differs in the position of dorsal spine on cerci which is present sub-basally in both *P. gravelyi* (small) and *P. mortoni* (long and prominent). In the new species, a small dorsal spine is present on the cerci after its forking, on the outer fork. Also, both *P. gravelyi* and *P. mortoni* are larger (AB+CA > 42 mm) with significantly distinct markings on prothorax. In *P. gravelyi*, prothorax is yellowish white with the broad black marking on the dorsum of its posterior lobe spreading to the middle lobe in a trapezoid fashion. In *P. mortoni*, prothorax is pale blue, except the posterior lobe which is black dorsally with a blue spot at the center. In the new species, prothorax is pale purple with the dorsum of posterior lobe marked with a broad, black, forward-curving stripe that extends into the middle lobe as two ‘ears’. The genital ligula among the three species also exhibits differences, as depicted (Figure 8). The forked tips of the third segment are rounded in the new species, while they are shovel shaped in *P. gravelyi* and end as curved hooks in *P. mortoni*.

**Figure 5:**
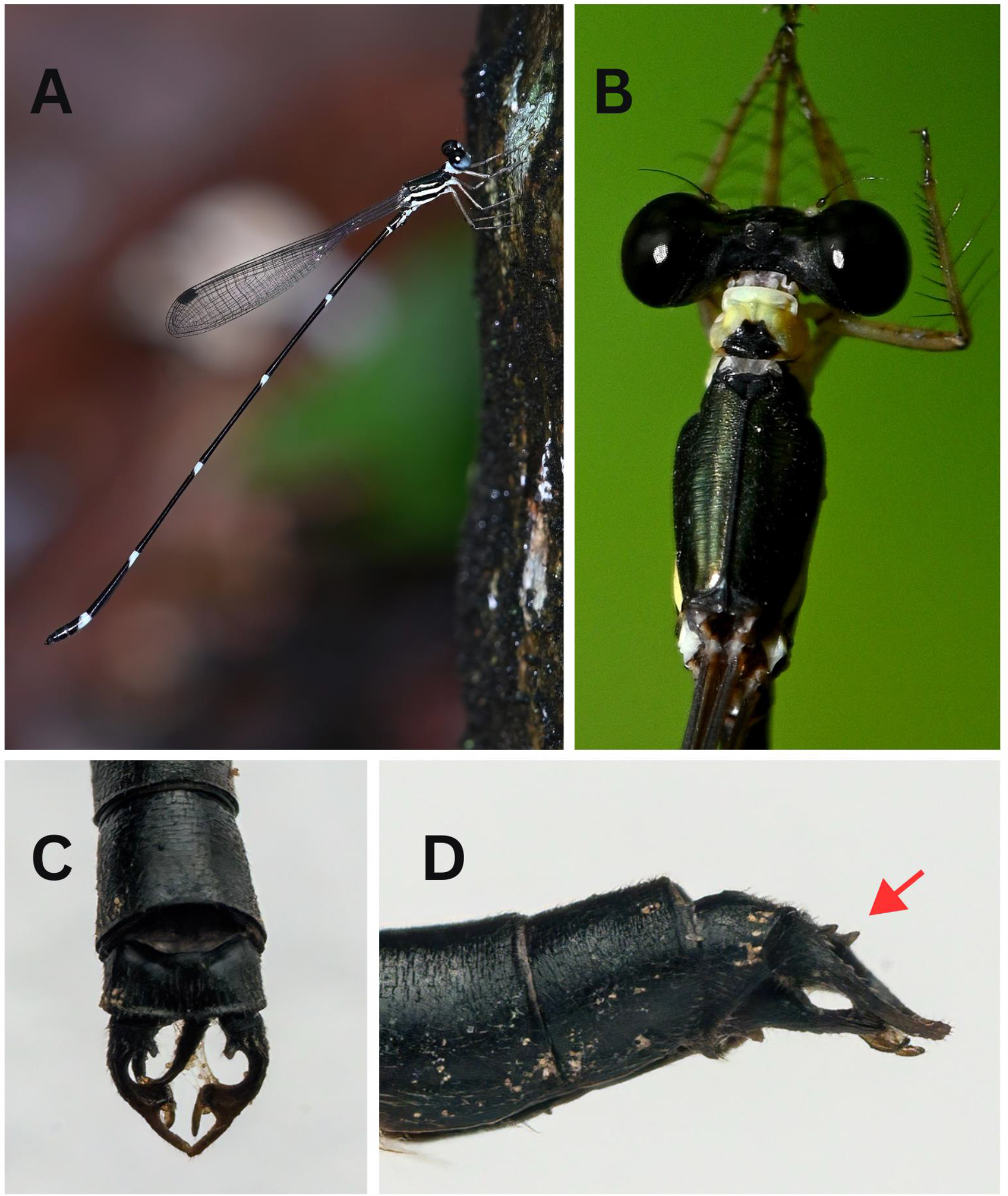
Male *Protosticta gravelyi*, A- in the habitat, B- dorsal view of head and thorax, C- dorsal view of caudal appendages, D- lateral view of caudal appendages.

**Figure 6:**
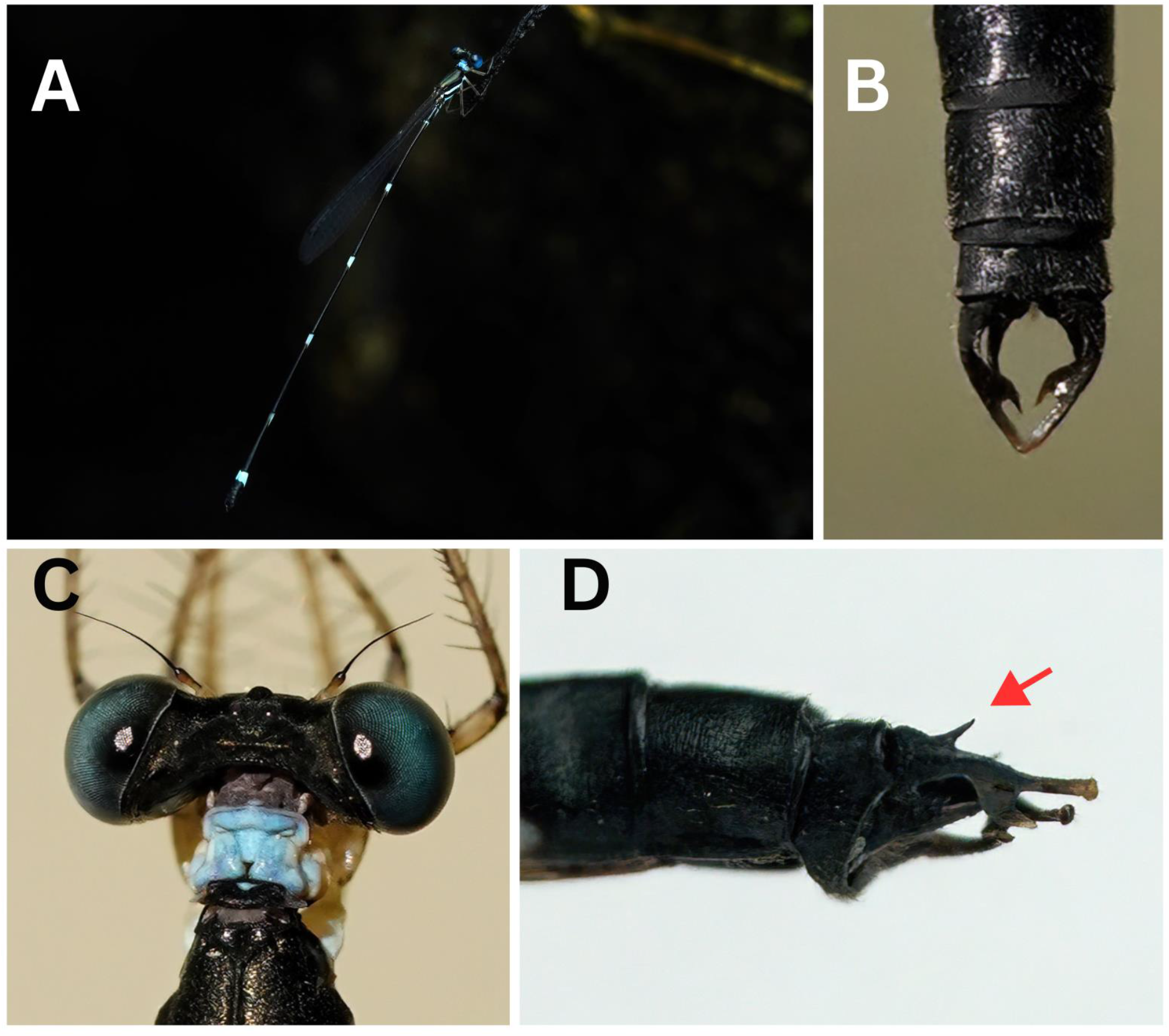
Male *Protosticta mortoni*, A- in the habitat, B- dorsal view of caudal appendages, C- dorsal view of head and prothorax, D- lateral view of caudal appendages.

**Figure 7:**
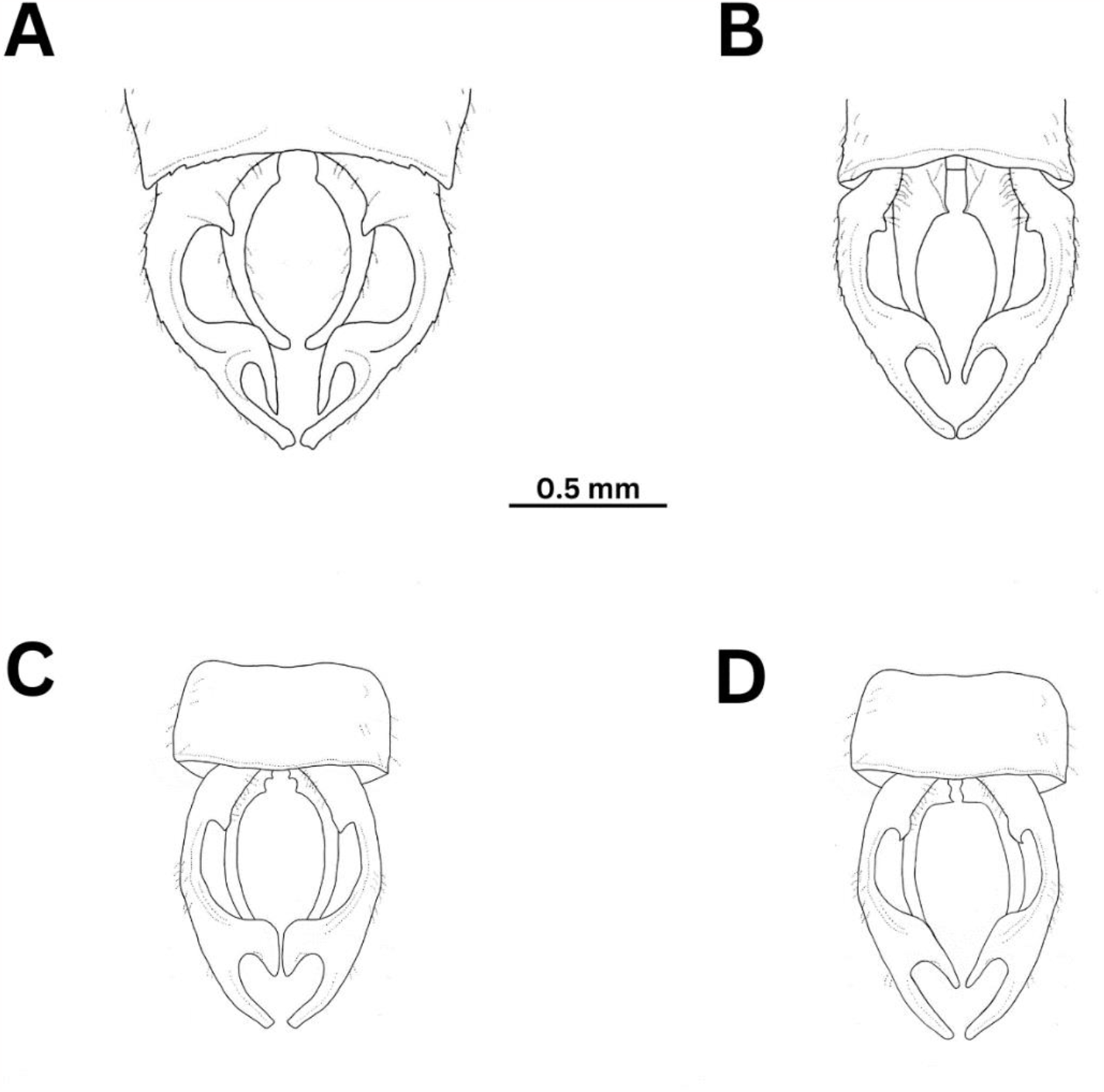
Illustrations of dorsal view of caudal appendages of: A- *Protosticta gravelyi*, B- *P. mortoni*, C- *P. hearseyi*, D- *P. sexcoloratus* sp. nov.

**Figure 8:**
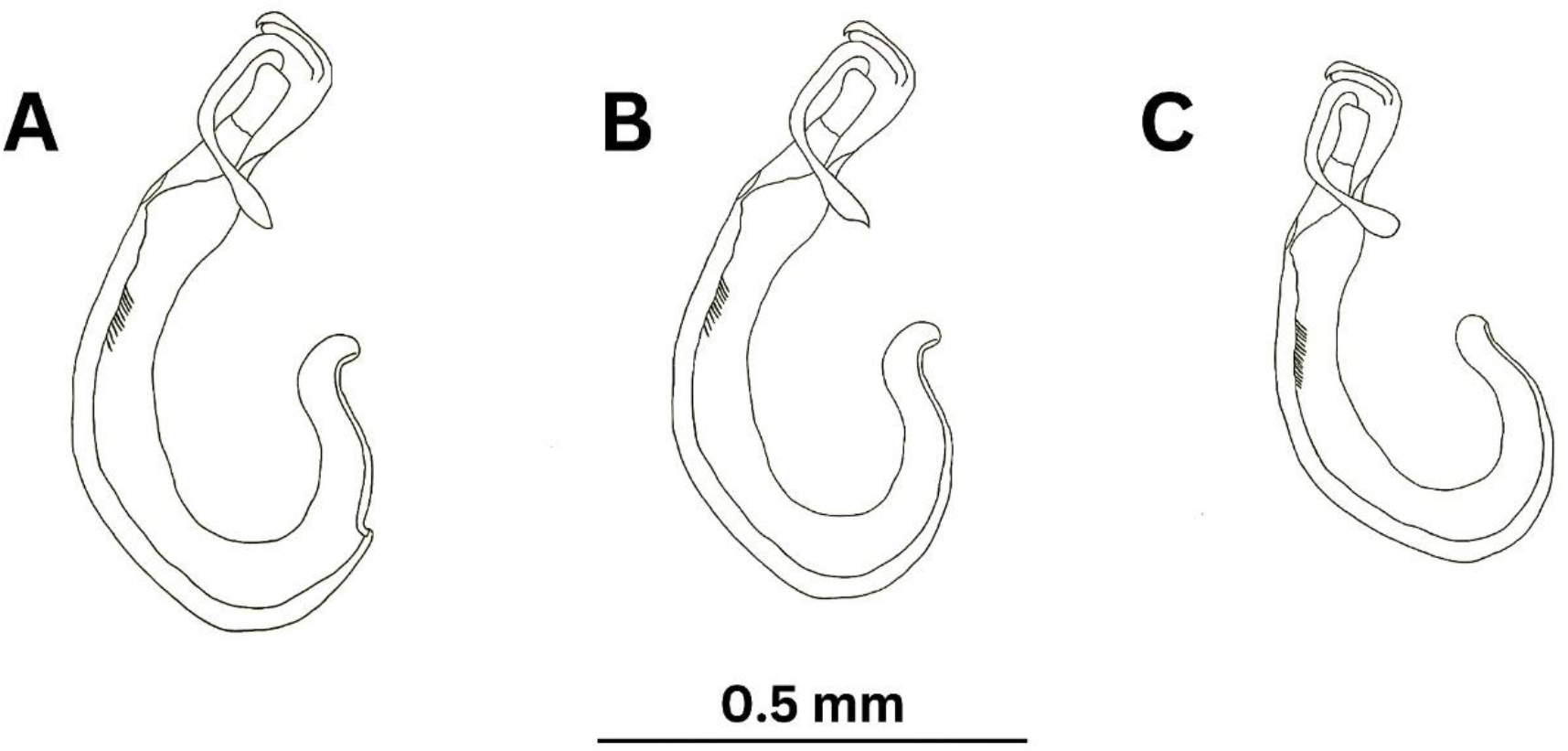
Illustrations of genital ligulae of: A- *Protosticta gravelyi*, B- *P. mortoni*, C- *P. sexcoloratus* sp. nov.

*Protosticta davenporti* Fraser, 1931 and *P. rufostigma* Kimmins, 1958 are also larger than the new species (AB+CA > 42 mm) and have pale yellow anterior and middle lobes of prothorax, the latter with a fine black mark spreading from the fully black posterior lobe. In both these species, paraprocts lack the basal spine.

The recently described *Protosticta sholai* Subramanian & Babu, 2020 is larger (AB+CA = 42 mm) and has S9 marked laterally with yellow. The tips of cerci are squared in this species. In the new species, S9 is unmarked and tips of cerci are rounded. *P. monticola* Emiliyamma & Palot, 2016 is about the same size as the new species, but has a brownish white prothorax marked with black, S1 and S2 bright yellow laterally and ventrally, and S8 fully black dorsally. The new species has pale purple prothorax marked with black, S1 and S2 bluish white laterally and ventrally, and apical half of S8 bluish white dorsally.

*Protosticta sanguinostigma* Fraser, 1922 is larger (AB+CA > 42 mm), has a reddish-brown equatorial band in its bottle-green eyes, blood-red pterostigma, and cerci with long, pointed spines. The new species has bright blue eyes capped with dark brown, dark brown pterostigma, and cerci with short, inward curving, sub-basal spines. The recently described *P. myristicaensis* is the smallest Indian *Protosticta* (AB+CA < 20 mm) with turquoise blue prothorax and cerci with a tubercle in its apical fork. Compared to it, the new species is larger and can be told apart by its pale purple prothorax marked with black and absence of any tubercle on its cerci.

*Protosticta cyanofemora* Joshi, Subramanian, Babu & Kunte, 2020 is a closely similar species described recently from the Western Ghats, but the ventral side of its thorax is creamy white with a ‘Y’- shaped marking (thorax pale white ventrally, dark between the legs, and with a black stripe extending shortly after the hindlegs in *Protosticta sexcoloratus* sp.nov.). Also, in *P. cyanofemora*, the forking of cerci is not deep and the inner fork is short (forking of cerci deep in *P. sexcoloratus* sp.nov. with longer inner fork). Further, the genital ligula of *P. cyanofemora* is without any setae, but the first segment of genital ligula of *P. sexcoloratus* sp. nov. has 16 setae.

The new species can be easily told apart from both *Protosticta anamalaica* Sadasivan, Nair & Samuel, 2022 and *P. armageddonia* Chandran, Payra, Deshpande & Koparde 2023, both of which have brownish prothorax and shallow forking of cerci (prothorax pale purple marked with black and deep forking of cerci in the new species).

*Description of holotype*. (Figure 9). Male. TL 40 mm; AB+CA 34.5 mm; HW 20 mm.

**Figure 9:**
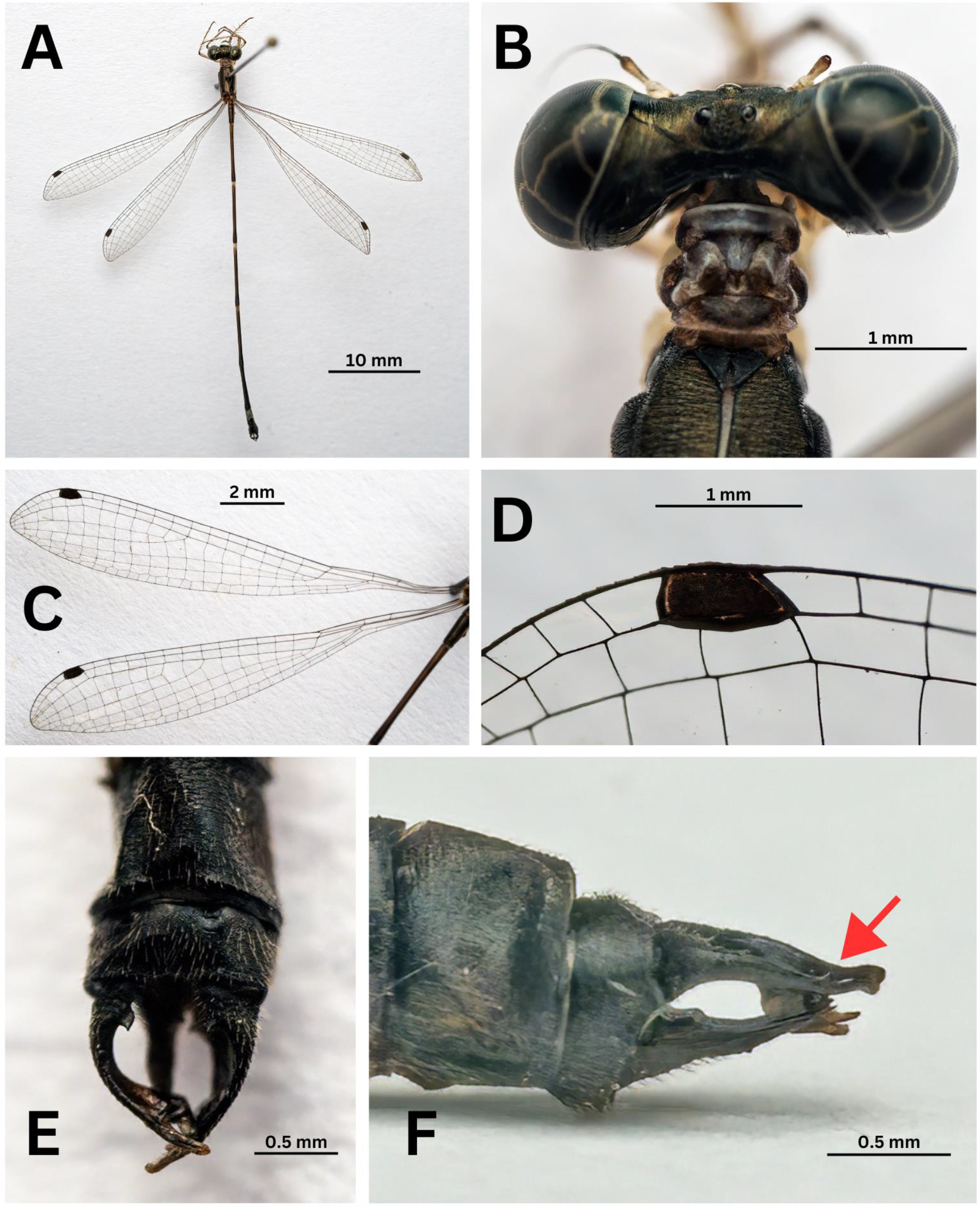
*Protosticta sexcoloratus* sp. nov. holotype male. A- habitus, B- prothorax, C- wings, D- pterostigma, E- caudal appendages in dorsal view, F- caudal appendages in lateral view.

Head. Labium white, tipped with dark brown; labrum bluish white with broad, black anterior margin; anteclypeus bluish white; bases of mandibles black; postclypeus, antefrons, postfrons and vertex steely black; ocelli dark brown; occiput black; bases of antennae bluish white, filaments dark brown; eyes bright turquoise blue anteriorly, antero-dorsally and postero-laterally, dark brown postero- dorsally.

Prothorax. Pale purple marked with black; anterior lobe with black, irregular inner margin; lateral lobes of middle lobe with large black spots; posterior lobe with a wavy outer margin, large, black forward curving mark medially, and pale purple collar; propleuron pale white at the coxal end, broadly black in the middle and pale purple at top.

Pterothorax. Black marked with bluish white; middorsal carina fine blue; mesostigmal plate, mesepisternum and mesepimeron steely black; mesinfraepisternum black; metepisternum and metepimeron bluish white; a broad black stripe over the metapleural suture; ventrally black between the legs, rest pale white with a black stripe extending shortly beyond the hindlegs.

Legs. Bluish white and black; coxae and trochanters bluish white; femora bluish white with black lines on extensor surface; knees black; tibia pale bluish white enfumed with black, apical half fully black; tarsi and tarsal claws black.

Wings. Hyaline; Pt dark brown, one cell-long, braced between thick, black nervures; Ab absent; Ax 2 in all wings; Px 13 in FW and 12 in HW; one cell between the junction of RP1–RP2 and the origin of IR2 in FW and HW.

Abdomen. Blackish brown marked with pale blue; S1 and S2 pale blue laterally; S3 to S7 with narrow pale blue basal annules extending more broadly along the sides and ventrally, fading in S6 and almost inconspicuous on S7; S8 turquoise blue, with a narrow black apical annule; S9 and S10 unmarked. S7 more than double the length of S8, S8 broad, double the length of S9, and S10 half the length of S9.

Caudal appendages. Black with brown tips; cerci about twice the length of S10, broad at the base and furnished here with a robust, inwardly directed sub-basal spine, then constricted and forked at the apical third, both outer fork and inner fork finger-like and rounded at the tip, the inner fork falling shorter and outer fork with a small, dorsal spine, the whole appendage curving gently in and downwards; paraprocts about four-fifths the length of cerci, with a blunt sub-basal tooth, broad at base, tapering to a spatulate, upturned tip.

*Paratype* 1. Male. TL 39.5 mm; AB+CA 34 mm; HW 19.5 mm.

Px 13 in both FW, 12 in left HW and 13 in right HW; rest agreeing with holotype.

Genital ligula. As illustrated in Figure 8C, first segment broad and curved, with 16 setae towards its end; second segment short and cylindrical; tip of third segment produced into two transparent filaments ending in rounded, club-like tips.

*Paratype* 2. Male. TL 40 mm; AB+CA 34 mm; HW 20 mm.

Px 13 in left FW, 12 in right FW, 12 in left HW and 11 in right HW; rest agreeing with holotype.

In this specimen, tips of caudal appendages of one side were broken. This allowed for a more detailed view of its complex structure (Figure 10).

**Figure 10:**
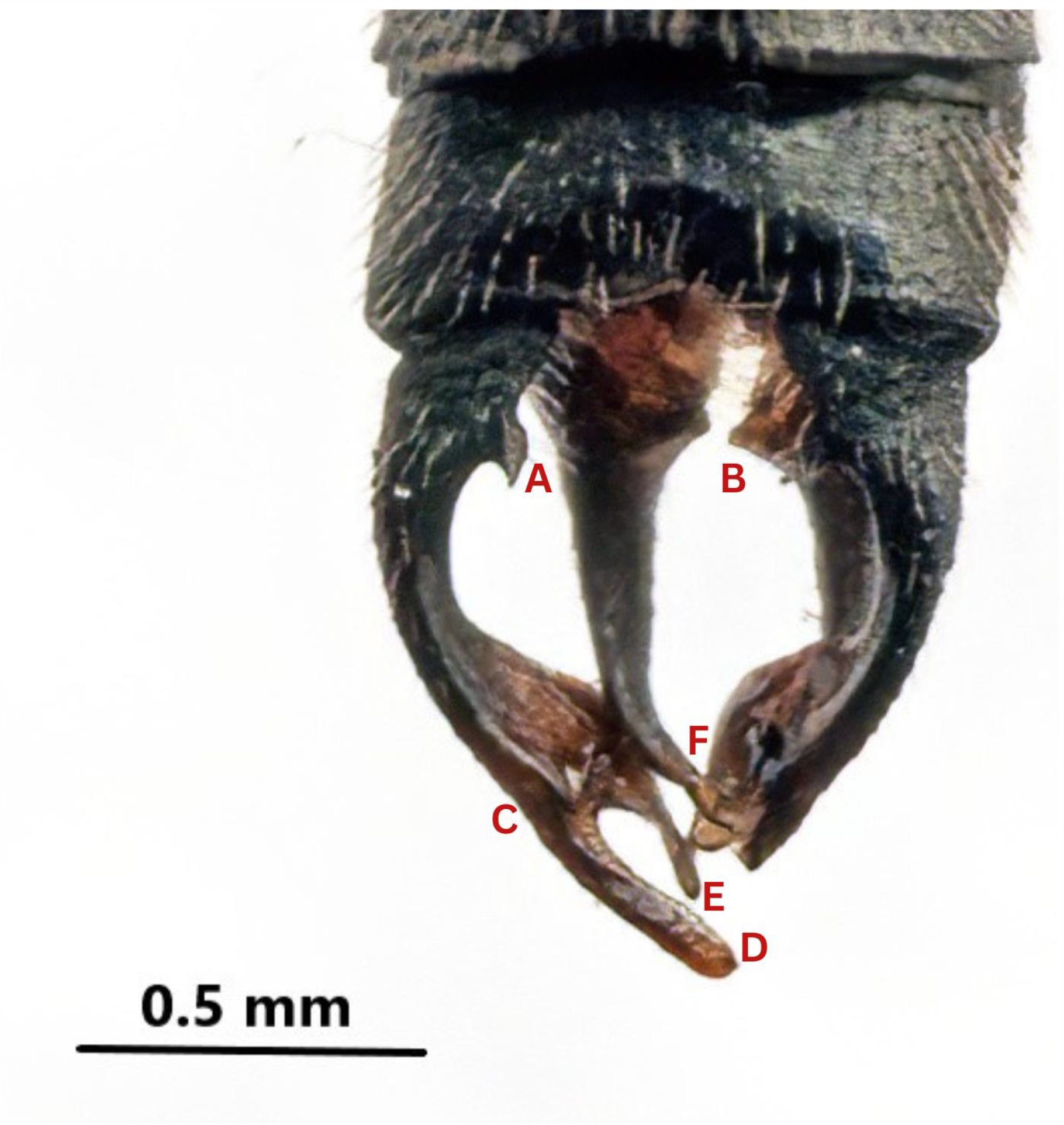
Dorsal view of *Protosticta sexcoloratus* sp. nov. paratype male. A- sub-basal spine of cerci, B- blunt sub-basal tooth of paraproct, C- small dorsal spine on cerci, D- rounded tip of cerci, E- long inner fork of cerci with rounded tip, F- spatulate tip of paraproct.

*Paratype* 3. (Figure 11). Female. TL 40 mm; AB+CA 33.5 mm; HW 22 mm.

**Figure 11:**
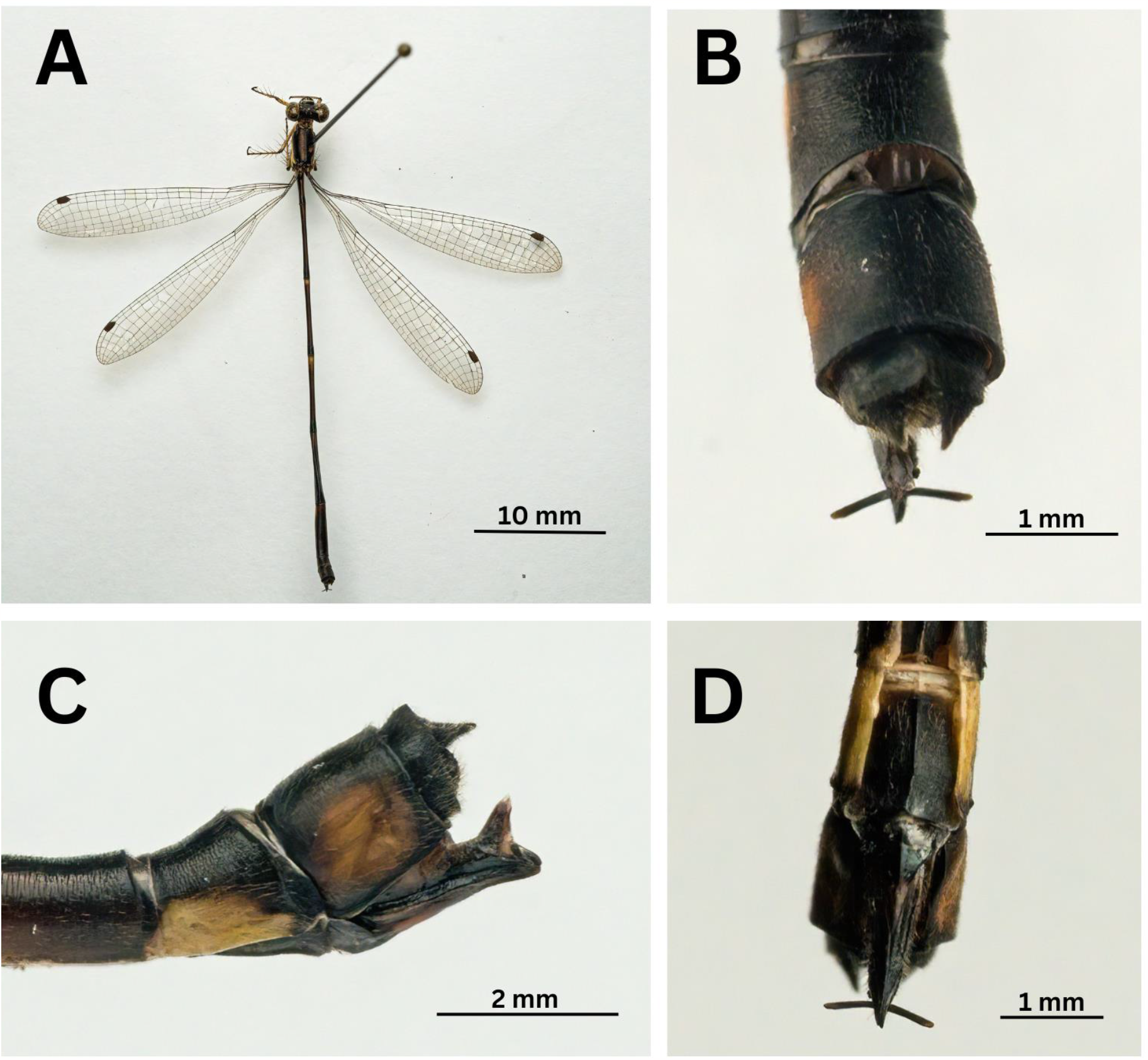
*Protosticta sexcoloratus* sp. nov. paratype female. A- habitus, B- dorsal view of last abdominal segments, C- lateral view of last abdominal segments, D- ventral view of last abdominal segments.

Similar to the male, but with the following differences: eyes apple green anteriorly, antero-dorsally and postero-laterally, reddish brown postero-dorsally; middorsal carina black (not marked in pale blue); pale purple of prothorax in male replaced with yellowish brown; pale blue/bluish white of thoracic markings, legs and abdominal markings in male replaced with pale yellow; S8 black dorsally with pale yellow marking extending to three-fourth of the segment ventro-laterally; S9 with broad yellowish brown mark laterally; S7 thrice the length of S8; S8 and S9 of about same length, but S9 broader; S10 less than half the length of S9.

Wings. Pt of HW longer and narrower than of FW; Ax 2 in all wings; Px 14 in left FW, 13 in right FW, 14 in both HW.

Caudal appendages. Black; covered with fine brown hairs; cerci short, as long as S10, conical with pointed ends; vulvar scale robust, black, with a yellow margin at base; styli black and long with paler ends; ovipositor extends beyond the abdomen.

*Paratype* 4. Female. TL 39 mm; AB+CA 33 mm; HW 21 mm.

Px 14 in all wings; rest agreeing with the former type.

*Remarks*. Many *Protosticta* species are closely similar to each other and require careful examination for identification. The new species is most similar to *P. hearseyi*, but can be distinguished as per the characters discussed in diagnosis. In this study, we have used the images of lectotype of *P. hearseyi* (British Natural History Museum 2023) for comparative analysis.

*Etymology*. The species epithet *sexcoloratus* highlights the difference in coloration between the sexes. In most *Protosticta* species, the male and female do not differ starkly in colors, the latter being a shorter and stouter version of the former. In *Protostcita sexcoloratus* sp. nov. however, the sexes are of the same length, but show distinct difference in coloration.

*Distribution*. Presently, the species has been recorded only from Vellarimala and 900 Kandi (11.502570° N, 76.110477° E, 1200 m above MSL), both of them peaks of the Camel’s Hump Mountains of Wayanad, a part of the Southern Western Ghats. Considering the known distribution of *Protosticta* species of the Western Ghats, we can assume that the new species is restricted to the high-altitude peaks of the Camel’s Hump Mountains.

*Temporal distribution*. Recorded in May (12) and June (3).

*Natural history*. All the specimens in this study were collected from shrubbery at the sides of precipitous ravines in evergreen jungle, near small, seasonal streams. Males and females were seen in about equal numbers, perched on vegetation not more than 2 meters above the ground.

*Discussion*. This study adds the 16^th^ species of *Protosticta* to the Western Ghats, seven of which were described after 2020. Taking into account the ecology of these species, we can surmise that there could be more *Protosticta* species awaiting discovery in the Western Ghats. Many of the *Protosticta* species have narrow distribution ranges within this biodiversity hotspot, emphasizing the need for detailed ecological studies and conservation initiatives focused on them. Dedicated explorations of other peaks in the Western Ghats, especially during the monsoon months, could result in more species discoveries from this group. As inventorying serves as the initial step in the process of biodiversity conservation, authorities and funding agencies should prioritize supporting such expeditions to document taxa hitherto unknown to science.

## Acknowledgements

We convey our sincere thanks to the Kerala Forests & Wildlife Department, Government of Kerala, for permitting us to conduct this study and providing us with all logistic support (Collection permit Ref: KFDHQ-5492/2021-CWW/WL10 dt 13.04.2022). We thank the Principal, Christ College (Autonomous), Irinjalakuda, for facilitating the laboratory work. We gratefully acknowledge the help of Niranjana KS for the illustrations. Many thanks to Reji Chandran for permitting to use his photograph of *Protosticta gravelyi*. The first, second and third authors are grateful to Society for Odonate Studies (SOS) for the encouragement to study Odonata.

